# The Cancer/Testis Antigen FATE1 Antagonizes Fission and Preserves Mitochondrial Network Integrity under Cytotoxic Stress

**DOI:** 10.1101/2025.08.18.670752

**Authors:** Verena Lang, Jasmin Tina Zahn, Deepti Pai Tonse, Nathan Ryan Brady, Anne Hamacher-Brady

**Affiliations:** Lysosomal Systems Biology, German Cancer Research Center, Heidelberg, Germany; Center for Quantitative Analysis of Molecular and Cellular Biosystems (BioQuant), Heidelberg University, Heidelberg, Germany; Optical Nanoscopy Division, German Cancer Research Center, Heidelberg, Germany; BioQuant, Heidelberg University, Heidelberg, Germany; Institute of Biochemistry and Molecular Biology I, Medical Faculty and University Hospital Düsseldorf, Heinrich Heine University Düsseldorf, Düsseldorf, Germany; Department of Molecular Microbiology & Immunology, Johns Hopkins University, Bloomberg School of Public Health, Baltimore, Maryland 21205, USA

**Keywords:** Apoptosis resistance, cancer biology, cancer/testis antigen FATE1, Drp1, Mff, mitochondria, mitochondrial dynamics, Mitofusin-2 (Mfn2), programmed cell death, stress response

## Abstract

FATE1 (Fetal and Adult Testis Expressed 1), also known as BJ-HCC-2, is a cancer-testis antigen with highly restricted expression in normal tissues but aberrant activation in diverse tumor types, where it localizes to both the outer mitochondrial membrane and the endoplasmic reticulum (ER) and functions as a key survival factor. Although FATE1 shares sequence homology with the mitochondrial fission factor Mff, its role in regulating mitochondrial architecture has not been mechanistically defined. Here, we identify FATE1 as a novel modulator of mitochondrial morphology that acts through a mechanism distinct from Mff. Unlike Mff, FATE1 does not recruit Drp1 to mitochondria and therefore lacks canonical fission activity. Instead, FATE1 promotes mitochondrial hyperfusion and protects against mitochondrial fragmentation triggered by cytokines and mitochondrial uncoupler treatment. The pro-fusion activity of FATE1 requires its mitochondrial targeting and interaction with Mitofusin-2 (Mfn2). Our findings establish FATE1 as a cancer-selective regulator of mitochondrial dynamics that antagonizes fission and preserves network integrity under cytotoxic stress, revealing a potential mechanism by which tumor cells evade mitochondria-driven apoptotic signaling.

## INTRODUCTION

Mitochondria are dynamic organelles whose morphology and function are tightly linked to cellular stress responses and cell fate decisions (1, 2). For instance, morphological remodeling of the mitochondrial network, including via fission and fusion, plays a critical role in modulating processes such as apoptosis (3, 4) and mitophagy (5, 6). Among the core regulators of mitochondrial fission is the outer mitochondrial membrane (OMM) protein mitochondrial fission factor (Mff), which recruits the dynamin-related GTPase Drp1 to initiate mitochondrial division (7, 8). While a central role of Drp1 in promoting mitochondrial fragmentation during stress and apoptosis is well established (9–11), growing evidence points to a complex regulatory landscape. For instance, the mitochondrial elongation factors MiD49 (MIEF2) and MiD51 (MIEF1) can recruit Drp1 to mitochondria independently of Mff (12, 13). Intriguingly, heterologous overexpression of MiD49 or MiD51 leads to Drp1 accumulation at the OMM but paradoxically induces mitochondrial elongation, not fission, suggesting that Drp1 recruitment alone is insufficient to drive division (13). Mild upregulation of MiD49/51 expression on the other hand enhances mitochondrial fragmentation by promoting Drp1 oligomerization and fission foci formation (13). These dual, context-dependent roles underscore the imporance of regulatory proteins that fine-tune mitochondrial dynamics beyond canonical fission/fusion paradigms.

Cancer cells frequently exhibit profound alterations in mitochondrial dynamics, metabolism and apoptosis regulation, which contribute to their ability to evade cell death and adapt to metabolic stress (14–16). Given the tight coupling between mitochondrial morphology and cellular apoptosis sensitivity, context- or cell type-specific regulators of mitochondrial dynamics are of particular interest as potential modulators of therapy resistance and as candidate targets for cancer-selective intervention strategies. Identifying such factors, especially those aberrantly expressed in cancer, may offer new opportunities to disrupt survival pathways that are uniquely exploited by malignant cells. Fetal and adult testis expressed (FATE1), also known as BJ-HCC-2, is a cancer/testis (CT) antigen with highly restricted expression in normal tissues (17), but aberrant activation in a variety of tumor types, including hepatocellular, lung, gastric, colon and adrenal carcinomas (18–20). Notably, FATE1 shares significant C-terminal homology with Mff, including a predicted coiled-coil and transmembrane domain (7, 20), raising the possibility that it may play a role in mitochondrial dynamics. However, to date, it has not been established whether FATE1 regulates mitochondrial morphology, nor whether such a function would involve mechanisms similar to canonical fission factors like Mff.

Initial studies identified FATE1 as a cancer survival factor that confers resistance to pro-apoptotic stress. In a genome-wide siRNA screen, Whitehurst et al. identified the *FATE1* gene as a target that reduced small-cell lung cancer cell viability to otherwise sublethal concentrations of paclitaxel (21). Maxfield et al. demonstrated that FATE1 promotes degradation of the pro-apoptotic BH3-only protein BIK via the E3 ligase RNF183, thereby suppressing intrinsic apoptosis in multiple cancer types, including melanoma, ovarian, breast, prostate and lung cancers (20). Independently, Doghman-Bouguerra et al. showed that FATE1 localizes to mitochondria-associated membranes (MAMs), where it interacts with the endoplasmic reticulum (ER)-resident protein Emerin to increase ER-mitochondria distance, limiting calcium transfer and reducing apoptosis sensitivity in adrenocortical carcinoma cells (22). Interestingly, the authors observed that induction of FATE1 expression in adrenocortical carcinoma cells correlated with decreased mitochondrial fragmentation, though without mechanistic investigation into how FATE1 might regulate mitochondrial morphology. More recently, Gallegos et al. reported that FATE1 supports survival and anchorage-independent growth in Ewing sarcoma cells, potentially by targeting of the atypical BH3-only protein and mitophagy receptor Bnip3L/Nix for degradation (23). This builds on earlier work from our group and others implicating Bnip3 and Nix as dual-function BH3-only proteins that link apoptosis and mitophagy (24–28).

Despite the growing recognition of FATE1 as a cancer cell survival factor and potential therapeutic target, its molecular functions remain incompletely understood, particularly regarding its potential involvement in the regulation of mitochondrial morphology and dynamics. In this study, we systematically dissect the structural features and molecular interactions that govern FATE1’s mitochondrial localization and function, and investigate its impact on mitochondrial network organization. Our findings establish FATE1 as a novel, cancer cell-specific regulator of mitochondrial dynamics that antagonizes canonical fission pathways, promotes hyperfusion, and preserves mitochondrial network integrity under cytotoxic stress.

## RESULTS

### FATE1 shares sequence similarity with Mff and localizes to both mitochondria and ER

Sequence analysis showed that FATE1 shares high homology with the C-terminal region of the mitochondrial fission factor Mff, including with the Mff coiled-coil (CC) domain and transmembrane (TM) segment (**Fig. 1a-c**; **Supplementary Fig. S1**), consistent with prior reports (7, 20, 22). COILS-based prediction algorithms (29, 30) confirmed a strong propensity for coiled-coil formation in this region, comparable to Mff and in contrast to negative controls such as Bnip3 (**Fig. 1b**). Importantly, our analysis further revealed that while Mff contains two conserved N-terminal repeat motifs (R1 and R2) essential for Drp1 recruitment (8, 31), FATE1 harbors only a sequence resembling R2 and lacks an R1 homolog (**Fig. 1c**; **Supplementary Fig. S1**), suggesting that FATE1 may functionally diverge from Mff by lacking key determinants required for canonical Drp1 engagement.

**Figure 1:**
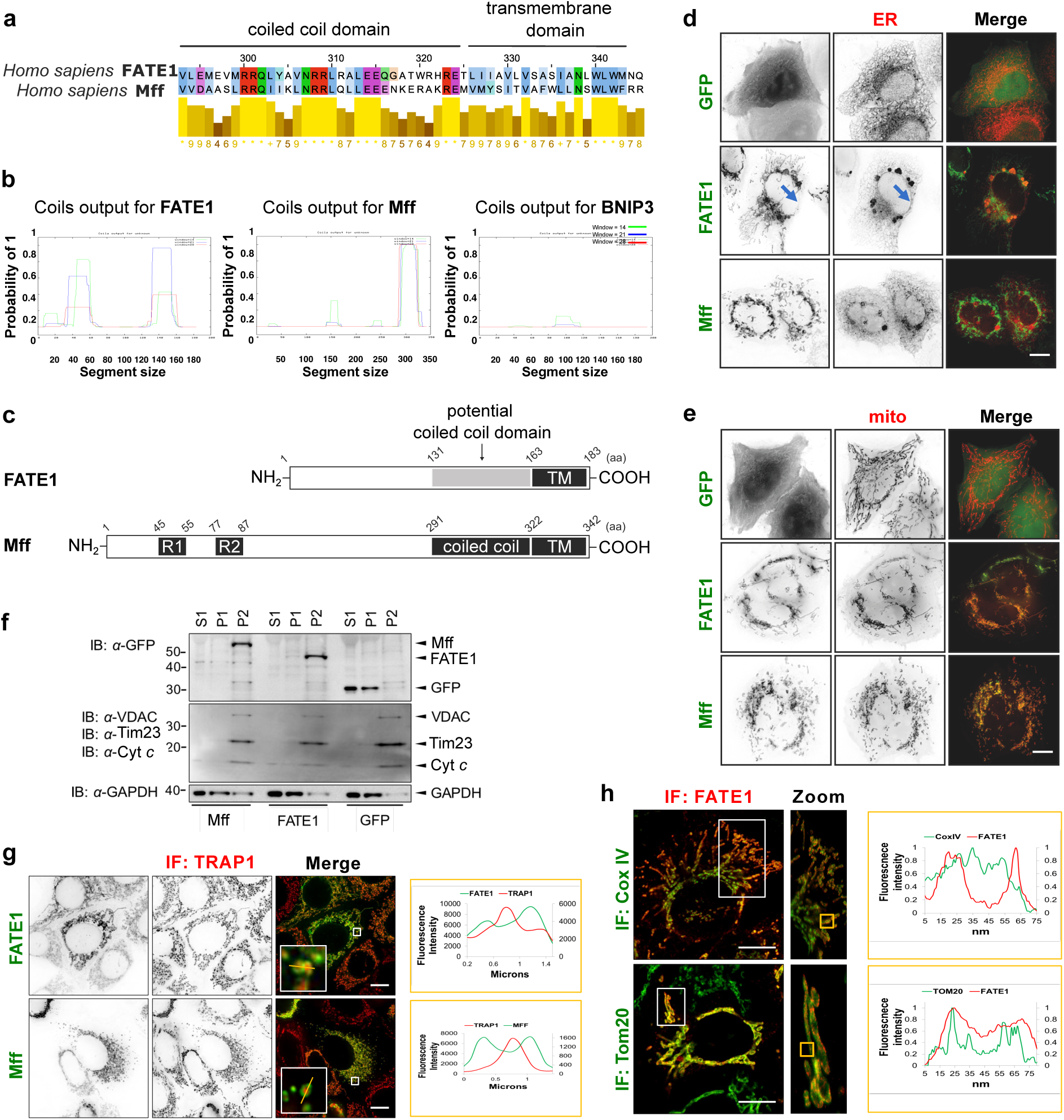
Sequence analysis and subcellular localization of FATE1 and Mff. **a** Sequence alignment of the C-terminal regions of human FATE1 and Mff generated with Jalview using the T-Coffee algorithm under default settings. Amino acid numbering corresponds to Mff. **b** Coiled coil domain prediction of FATE1 using the COILS program. Mff and Bnip3 were included as positive and negative controls, respectively. Predicted probabilities for coiled coil formation are shown for scanning windows of 14, 21 and 28 residues. **c** Schematic comparison of the domain architecture of Mff and FATE1, highlighting sequence homologies, including the predicted coiled coil domain and transmembrane region. R1 and R2 indicate the short repeat motifs in Mff involved in Drp1 recruitment. **d** Representative widefield images of GFP-Ctrl, GFP-FATE1, or GFP-Mff co-expressed with RFP-KDEL (ER marker) for 24 h in HeLa cells. Blue arrows, ER-localized FATE1 at perinuclear regions. **e** Representative widefield images of GFP-Ctrl, GFP-FATE1, or GFP-Mff co-expressed with mito-RFP (mitochondrial marker) for 24 h in HeLa cells. **f** Subcellular fractionation of HeLa cells transfected with GFP-tagged constructs as indicated. Fractionation yielded cytosolic (S1), rest-mitochondrial (P1) and purified mitochondrial (P2) fractions, analyzed by immunoblotting. **g** Representative widefield images of MCF-7 cells expressing GFP-FATE1 or GFP-Mff for 24 h and IF stained for mitochondrial matrix protein TRAP1. Plot profiles along yellow lines in zoom images show fluorescence intensity of GFP and TRAP1 signals. **h** STED super-resolution microscopy of HeLa cells expressing pcDNA-FATE1 and IF stained for FATE1 in combination with Cox IV (IMM marker) or Tom20 (OMM marker). Plot profiles show fluorescence intensity of FATE1 and Cox IV or Tom20. Scale bars, 10 µm

Next, we directly compared GFP-tagged FATE1 and Mff localization in HeLa cervical and MCF-7 breast cancer cells using organelle-specific markers, to validate and extend previous reports describing FATE1 localization to mitochondria and ER (20, 22). As expected, Mff localized exclusively to mito-RFP labeled mitochondria (**Fig. 1d-e**). In contrast, FATE1 was detected both at mitochondria and the ER, co-localizing with RFP-KDEL and exhibiting pronounced enrichment in perinuclear ER regions (**Fig. 1d**, blue arrows). Notably, FATE1 overexpression induced formation of puncate ER clusters not observed in control cells, suggesting the capacity for FATE1-driven ER remodeling. These findings confirm dual-organelle localization while providing side-by-side comparison to Mff and new evidence for morphological changes associated with FATE1 expression. Subcellular fractionation experiments further confirmed the dual localization of FATE1. GFP-FATE1 was enriched in the mitochondrial fraction (P2) and also detected in the light membrane fraction (P1), which contains ER-derived membranes, consistent with its localization to both mitochondria and the ER (**Fig. 1f**). In contrast, GFP-Mff was exclusively present in the mitochondrial fraction (P2), reflecting its selective mitochondrial targeting. GFP-Ctrl, as expected, was distributed across the cytosolic (S1) and light membrane (P1) fractions.

High-resolution widefield (**Fig. 1g**) and super-resolution STED (**Fig. 1h**) microscopy further clarified the submitochondrial localization of FATE1. In particular, zoom images and plot profile analyses of fluorescence intensities along a line crossing through single mitochondria revealed that FATE1, similar to Mff, circumscribed inner mitochondrial compartments labeled with the matrix localized chaperone TRAP1 (**Fig. 1g**). Similarly, STED microscopy zoom images and plot profile analyses showed that FATE1 surrounded IMM marker Cox IV, while colocalizing with OMM marker Tom20. These data support that FATE1 predominantly localizes to the OMM.

In summary, FATE1 and Mff share structural motifs in their C-termini, including a coiled-coil and TM domain, but differ in subcellular localization. While Mff is restricted to mitochondria and contains two conserved N-terminal repeat motifs (R1 and R2), critical for Drp1 recruitment and mitochondrial fission, FATE1 exhibits dual localization to both the OMM and the ER, and contains only a motif resembling R2, with no R1 homolog identifiable. This structural divergence, together with FATE1’s distinct localization and induction of ER remodeling, suggests that despite the sequence homology, FATE1 is unlikely to act as a canonical fission adaptor like Mff and may fulfill unique, Drp1-independent roles in organelle dynamics and cellular homeostasis.

### FATE1 does not function as a mitochondrial Drp1 receptor

As a first functional inquiry into FATE1’s potential role in mitochondrial dynamics, we compared its ability to engage Drp1 with that of the canonical Drp1 adaptor Mff. Overexpression of Mff is sufficient to drive Drp1 relocation to mitochondria and promote mitochondrial fission, by recruiting Drp1 to the outer mitochondrial membrane via conserved N-terminal R1 and R2 repeat motifs (8, 31, 32). Although FATE1 shares partial sequence similarity with Mff, including a motif resembling R2, it lacks an R1 homolog (**Fig. 1a-c**; **Supplementary Fig. S1**), suggesting limited or absent capacity to recruit Drp1.

To test this, we performed co-immunoprecipitation assays in HeLa cells co-expressing GFP-tagged Mff or FATE1 with either RFP-tagged wild-type Drp1 (Drp1-wt) or dominant-negative, GTPase-deficient Drp1 mutant (Drp1-DN) (33, 34). As expected, Mff strongly co-immunoprecipitated with Drp1-wt and to a lesser extent with Drp1-DN (**Fig. 2a**), consistent with preferential binding to GTPase-active Drp1 oligomers (32). In contrast, FATE1 showed only weak association with either form of Drp1, and endogenous Drp1 co-immunoprecipitated with Mff but not with FATE1.

**Figure 2:**
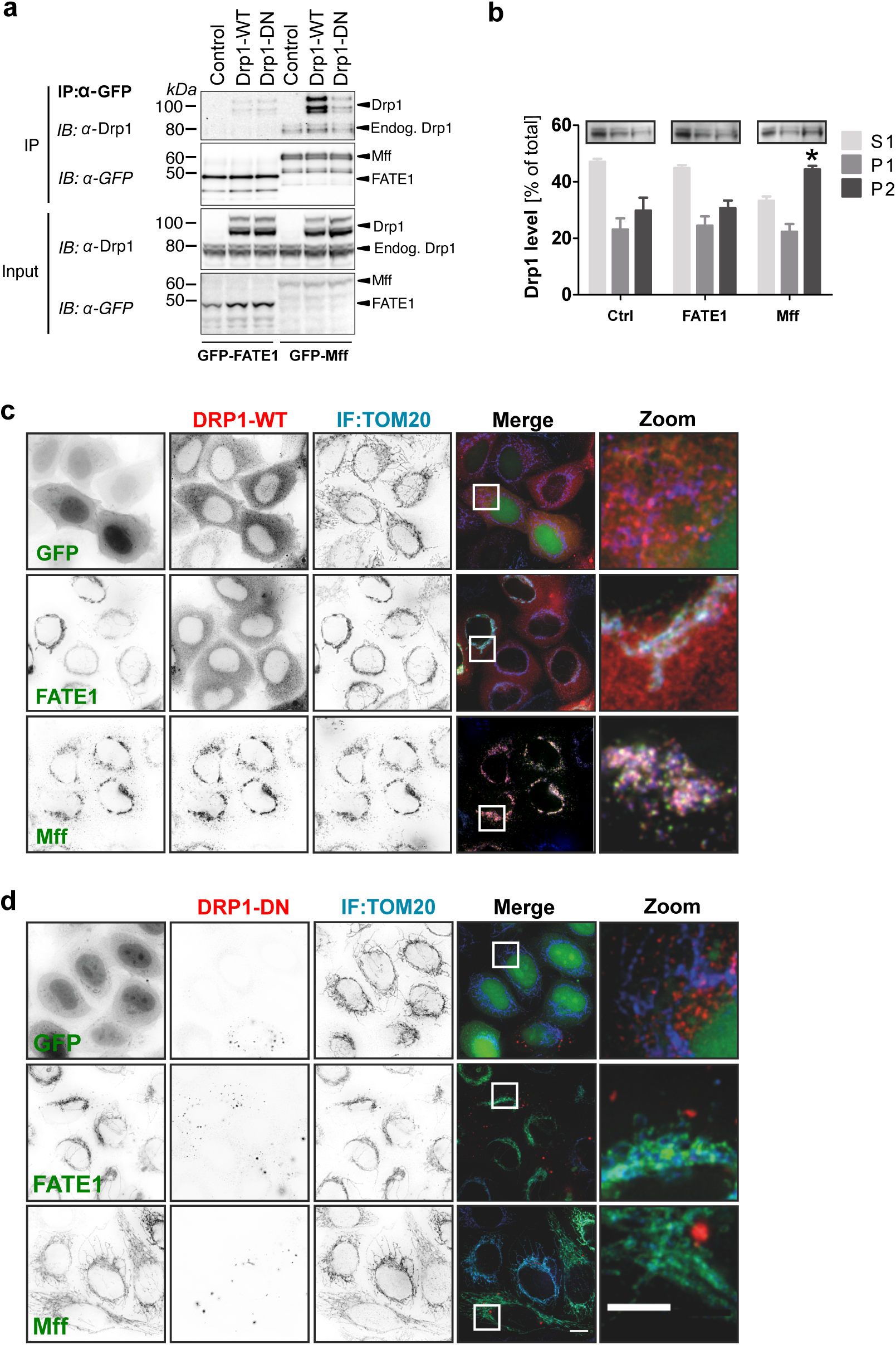
FATE1 does not recruit Drp1 to mitochondria or function as a Drp1 receptor. **a** Co-immunoprecipitation (co-IP) analysis of GFP-tagged FATE1 or Mff co-expressed with RFP-tagged Drp1-WT or Drp1-DN in HeLa cells. Cells were lysed 24 h post-transfection, and co-IP was performed using anti-GFP antibody. Immunoblots were probed with antibodies against Drp1 and GFP. **b** Subcellular fractionation of HeLa cells expressing GFP-Ctrl, GFP-FATE1 or GFP-Mff. Drp1 distribution was assessed across cytosolic (S1), rest-mitochondrial (P1), and mitochondrial (P2) fractions by immunoblotting. Quantification from three independent experiments is shown. Statistical analysis compared to GFP-Ctrl by one-tailed paired Student’s t-test; \**p* ≤ 0.05. **c** Representative widefield images of HeLa cells co-transfected with RFP-Drp1 and GFP-Ctrl, GFP-FATE1 or GFP-Mff. Mitochondria were visualized by IF staining for endogenous Tom20. **d** Representative widefield images of HeLa cells co-transfected with RFP-Drp1-DN and GFP-Ctrl, GFP-FATE1 or GFP-Mff. Mitochondria were visualized by IF staining for endogenous Tom20. Scale bars: overview, 10 µm; zoom, 5 µm.

Subcellular fractionation further demonstrated that Drp1 was enriched in mitochondrial fractions of Mff overexpressing cells, but remained largely cytosolic in FATE1 overexpressing cells (**Fig. 2b**). Consistently, fluorescence imaging of HeLa cells co-expressing either RFP-Drp1-wt or RFP-Drp1-DN together with GFP-FATE1, GFP-Mff, or GFP-Ctrl revealed that Mff, but not FATE1, strongly induced mitochondrial recruitment of Drp1-wt (**Fig. 2c**). Neither Mff nor FATE1 promoted substantial mitochondrial recruitment of Drp1-DN, although Drp1-DN expression potently inhibited Mff-induced mitochondrial fragmentation, confirming its dominant-negative function (**Fig. 2d**).

Together, these results demonstrate that FATE1, despite sharing partial sequence homology with Mff, does not act as a functional Drp1 receptor. It fails to bind or recruit Drp1 to mitochondria and lacks the sequence features required for assembly of Drp1 fission complexes at the outer mitochondrial membrane.

### FATE1 alters mitochondrial morphology and promotes hyperfusion

To determine whether FATE1 influences mitochondrial architecture independently of Drp1 recruitment, we systematically quantified morphological phenotypes in GFP-FATE1 expressing HeLa cells. Compared to control conditions, FATE1 overexpression markedly increased mitochondrial clustering (∼ 40%) and induced a distinct peri-nuclear clustering phenotype (∼ 10%) characterized by a single, condensed mitochondrial cluster adjacent to the nucleus (**Fig. 3a**). Morphometric analysis further revealed that FATE1 overexpression resulted in a shift from elongated to either fragmented or fused mitochondria, with ∼64% of cells exhibiting a non-fragmented phenotype (**Fig. 3b**). Notably, a substantial proportion (∼37.5%) of these non-fragmented cells displayed a hyperfused mitochondrial network, defined by highly interconnected tubules often spanning across or above the nucleus. The hyperfusion phenotype was especially evident in cells with high GFP-FATE1 expression levels.

**Figure 3:**
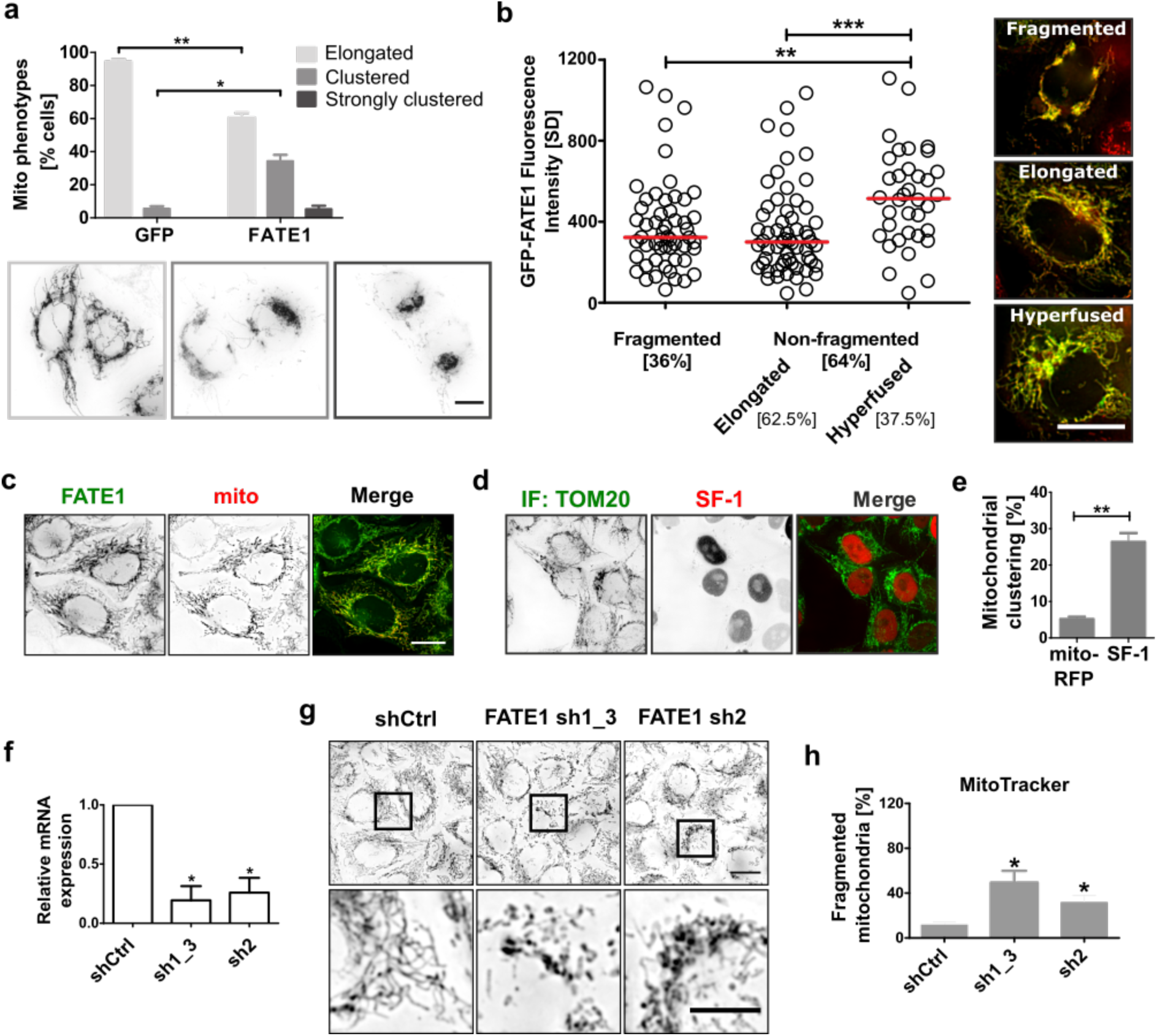
Enhanced FATE1 expression induces mitochondrial hyperfusion, while FATE1 depletion promotes fragmentation. **a** HeLa cells were transfected with OMM-GFP or GFP-FATE1, and mitochondrial morphology was assessed 24 h post-transfection. For each condition, 180 cells were analyzed by widefield microscopy per replicate in three independent experiments. Three major phenotypes were classified: elongated, clustered and peri-nuclear clustered mitochondria. **b** HeLa cells were co-transfected with GFP-FATE1 and mito-RFP to quantify mitochondrial hyperfusion within the non-fragmented population. Standard deviations of GFP-FATE1 pixel intensities (grey values) across mitochondrial regions were used to calculate mean variability per cell (red line), across a total of 150 cells (∼ 50 per experiment). Representative widefield images of the analyzed phenotypes are shown. Statistical comparisons were performed using Student’s unpaired t-test; \*\**p*<0.01, \*\*\**p*<0.001. **c** Representative widefield images of HeLa cells stably expressing GFP-FATE1 and transiently transfected with mito-RFP. **d** Representative widefield images of HeLa cells transiently transfected with RFP-SF-1 for 24 h. Tom20 IF was performed to visualize mitochondria. **e** Quantification of mitochondrial clustering from (**d**), expressed as the percentage of cells with clustered mitochondria. For each condition, 180 cells were analyzed in each of three independent experiments. Statistical comparisons to mito-RFP were made using paired Student’s t-test; \*\**p*<0.01. **f** RT-PCR analysis of FATE1 mRNA levels in HeLa cells following shRNA-mediated knockdown. Transcript levels were normalized to β-actin and are shown relative to shCtrl. Statistical significance was assessed using Student’s paired t-test; \**p*<0.05. **g** Representative widefield images of live shCtrl and shFATE1 HeLa cells stained with MitoTracker Red. **h** Quantification of non-fragmented vs. fragmented mitochondrial morphologies from (**f**), expressed as the percentage of cells with fragmented mitochondria. For each condition, 180 cells were analyzed in each of three independent experiments. Statistical comparisons to shCtrl were made using paired Student’s t-test; \**p*<0.05. Scale bars: overview, 10 µm; zoom, 5 µm.

To validate the link between FATE1 expression levels and mitochondrial elongation, we imaged stable GFP-FATE1 overexpressing HeLa cells, where uniformely lower fluorescence levels allowed clear visualization of mitochondrial morphology and at the same time revealed dual localization of FATE1 also to the ER (**Fig. 3c**). Moreover, overexpression of steroidogenic factor 1 (SF-1), a known transcriptional activator of FATE1 (35, 36), produced similar mitochondrial elongation and clustered phenotypes (**Fig. 3d-e**), indicating that SF-1 driven upregulation of endogenous FATE1 is sufficient to promote mitochondrial hyperfusion.

To enable loss-of-function analysis, we generated stable FATE1 deficient HeLa cells by lentiviral gene knockdown using three independent shRNAs targeting FATE1. Knockdown efficiency was confirmed at the RNA level (**Fig. 3f**). We observed that FATE1 depletion leads to a marked reduction in mitochondrial elongation phenotypes, as determined by MitoTracker Red staining (**Fig. 3g-h**), further supporting FATE1’s role in promoting mitochondrial elongation and network integrity.

In summary, FATE1 promotes mitochondrial elongation and hyperfusion, supporting mitochondrial network integrity and counteracting fragmentation. These findings suggest that FATE1 functions as a key modulator of mitochondrial morphology in cancer cells.

### Dissection of FATE1 domains controlling subcellular localization and mitochondrial morphology remodeling

To investigate how distinct structural domains of FATE1 contribute to its dual localization to mitochondria and the ER, as well as its ability to modulate mitochondrial morphology, we generated a panel of GFP-tagged deletion mutants (**Fig. 4a**). These mutants were designed to selectively disrupt the predicted coiled-coil (CC) and transmembrane (TM) domains (**Fig. 1a-c**; **Supplementary Fig. S1**), both of which have been implicated in mitochondrial localization of FATE1 in previous studies (20, 22). While prior work established the requirement of these domains for mitochondrial localization, their individual contributions to organelle-specific targeting remain incompletely defined, and their impact on morphological remodeling remains unknown. Here, we systematically assessed how loss of each domain affects FATE1’s subcellular localization and its capacity to drive mitochondrial elongation and hyperfusion.

**Figure 4:**
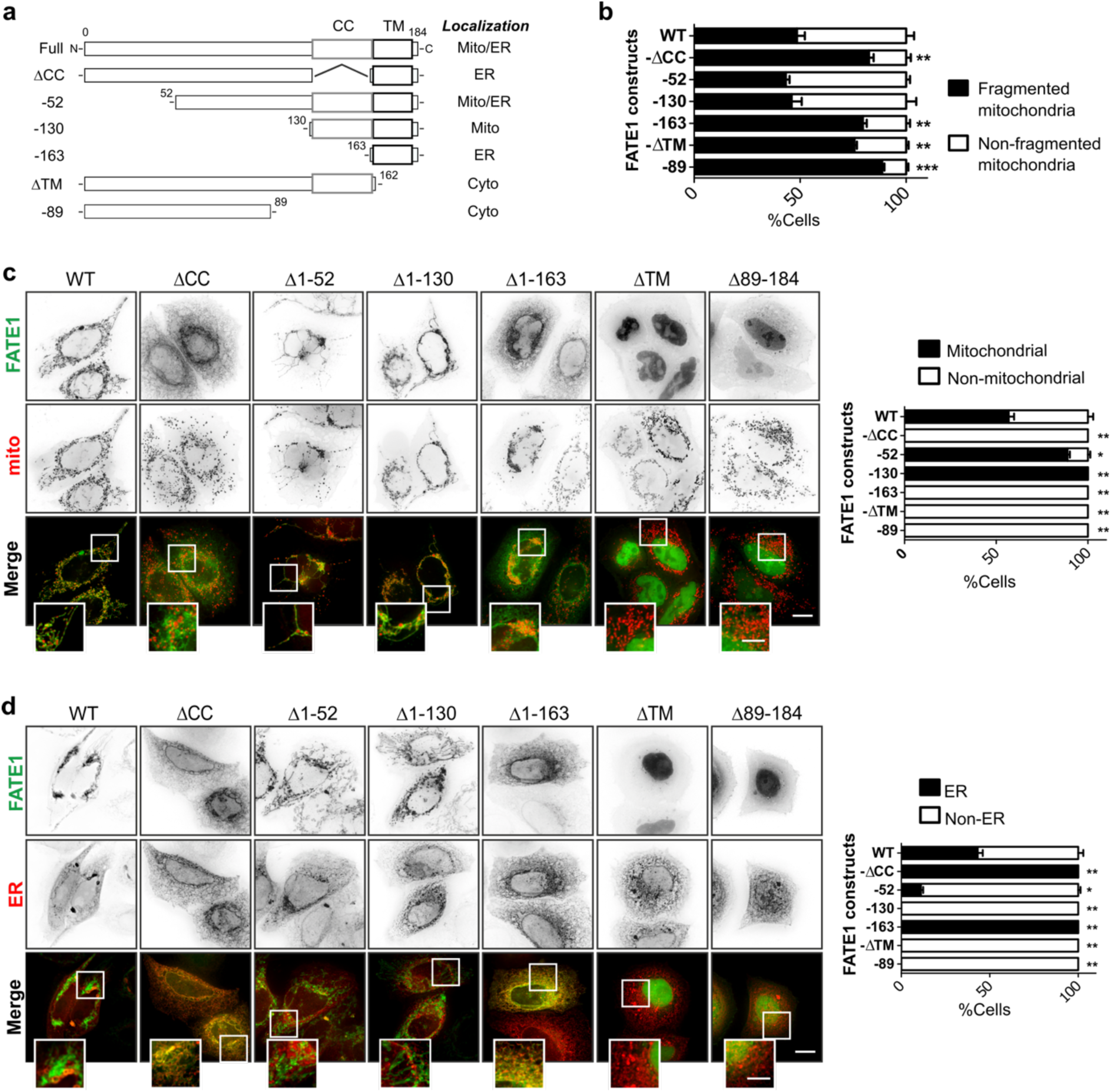
Subcellular localization and functional impact of FATE1 domain mutants. **a** Schematic overview of GFP-tagged FATE1 wild type (WT) and domain-deletion mutants, with summary of their observed subcellular localizations. **b** Quantification of mitochondrial fragmentation in HeLa cells expressing FATE1 mutants compared to WT. For each condition, 180 cells were analyzed in each of three independent experiments. Statistical comparisons were made to GFP-FATE1 WT using Student’s paired t-test; \**p*<0.05, \*\**p*<0.01, \*\*\**p*<0.001. **c** Representative widefield images of HeLa cells co-expressing GFP-FATE1 mutants with the mitochondrial marker mito-RFP. Quantification of mitochondrial vs. non-mitochondrial localization of FATE1 mutants. For each condition, 180 cells were analyzed in each of three independent experiments. Statistical comparisons were made to GFP-FATE1 WT using Student’s paired t-test; \**p*<0.05, \*\**p*<0.01. **d** Representative widefield images of HeLa cells co-expressing GFP-FATE1 mutants with the ER marker RFP-KDEL. Quantification of ER vs. non-ER localization of FATE1 mutants. For each condition, 180 cells were analyzed in each of three independent experiments. Statistical comparisons were made to GFP-FATE1 WT using Student’s paired t-test; \**p*<0.05, \*\**p*<0.01. Scale bars: overview, 10 µm; zoom, 5 µm.

Subcellular localization analysis revealed that the TM domain is essential for membrane integration of FATE1. A mutant lacking this region (ΔTM) was diffusely distributed throughout the cytoplasm and nucleus, phenocopying a comparable Mff mutant previously shown to mislocalize without its TM segment (31). Conversely, deletion of the predicted coiled coil domain (ΔCC) redirected FATE1 predominantly to the ER, indicating that this domain is critical for mitochondrial association but dispensable for ER localization (**Fig. 4b-e**). Similarly, a minimal TM-only mutant (residues 163-183) localized to the ER, consistent with its inability to target mitochondria in the absence of the CC domain. Simultaneous deletion of TM and CC (ΔTMΔCC) abolished membrane association entirely, resulting in diffuse cytoplasmic and nuclear distribution. In contrast, two N-terminal truncation mutants (Δ1-52 and Δ1-130), which retained both CC and TM domains, exhibited enhanced mitochondrial and reduced ER localization relative to full-length FATE1 (**Fig. 4a, 4c-d**). This suggests that regions within the N-terminus may negatively regulate mitochondrial targeting or enhance ER retention.

Functionally, the ability of FATE1 mutants to promote mitochondrial elongation and hyperfusion correlated with their subcellular localization. Morphometric quantification demonstrated that mutants failing to localize to mitochondria (ΔTM, ΔCC, ΔTMΔCC, TM-only) significantly incrased the fraction of cells with fragmented mitochondrial networks compared to wild-type FATE1 (**Fig. 4b**). In contrast, the mitochondrial localization-competent truncation mutants (Δ1-52 and Δ1-130) maintained non-fragmented mitochondrial morphology similar to FATE1-wt, further supporting a direct role of mitochondrial targeting in regulating FATE1’s pro-fusion activity.

Together, these findings delineate the distinct contributions of FATE1’s structural domains to its dual organelle localization and functional activity. The transmembrane domain is necessary for membrane integration, while the coiled coil domain mediates mitochondrial targeting.

### FATE1-induced mitochondrial hyperfusion is mediated by Mfn2

Above, we showed that FATE1-induced changes in mitochondrial morphology occur independently of the canonical fission factor Drp1, and identified a subpopulation of FATE1-expressing cells displaying a pronounced mitochondrial hyperfusion phenotype. To elucidate the molecular basis of FATE1-induced mitochondrial hyperfusion, we examined whether its effects depend on the mitochondrial fusion proteins, Mitofusin-1 (Mfn1) and Mitofusin-2 (Mfn2).

Co-immunoprecipitation assays in HeLa cells expressing GFP-FATE1 revealed a specific interaction with endogenous Mfn2, but not Mfn1 (**Fig. 5a**). Expression of FATE1 did not alter steady-state levels of Mfn1 or Mfn2 (**Fig. 5b**), supporting a model in which FATE1 modulates mitochondrial morphology through direct interaction with Mfn2 rather than via changes in mitofusin abundance.

**Figure 5:**
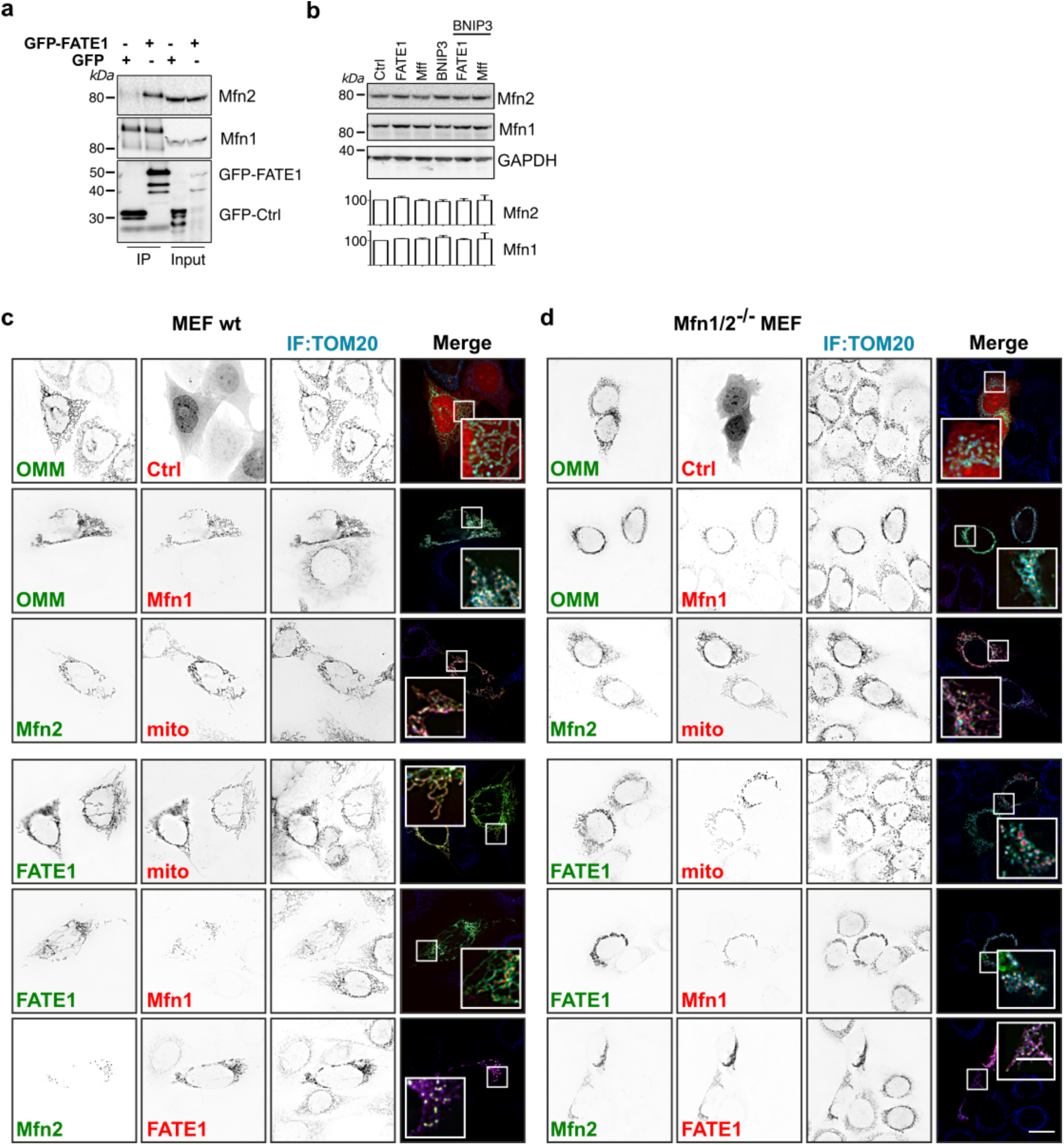
FATE1-induced mitochondrial hyperfusion is dependent on Mfn2. **a** Hela cells were transfected with GFP-Ctrl or GFP-FATE1 for 24 h, followed by co-immunoprecipitation using anti-GFP. Immunoblots were probed for GFP, and endogenous Mfn1 and Mfn2, to assess specific interactions. **b** Immunoblot analysis of endogenous Mfn1 and Mfn2 levels in HeLa cells transfected for 24 h with GFP-Ctrl, GFP-FATE1 or GFP-Mff, ± RFP-Bnip3. Immunoblots were probed for Mfn1 and Mfn2, and GAPDH as a loading control. Quantification from three independent experiments is shown. **c** Representative widefield images of wild type MEFs 24 h after transfection with mitochondrial markers OMM-GFP or mito-RFP in combination with GFP-Ctrl or RFP-Ctrl, YFP-Mfn2 or RFP-Mfn1, and FATE1 constructs (GFP- or RFP-tagged). Mitochondria were furthermore visualized by IF staining for endogenous Tom20. **d** Representative widefield images of Mfn1/2 DKO MEFs transfected and stained as in (**c**). Scale bars: overview, 10 µm; zoom, 5 µm.

To test the functional requirement of mitofusins in mediating FATE1’s effects, we expressed GFP-FATE1 in wild-type (wt) mouse embryonic fibroblasts (MEF) alongside mito-RFP, or co-expressed with YFP-tagged Mfn2 or RFP-Mfn1 (**Fig. 5c**). FATE1 overexpression alone induced profound mitochondrial elongation and clustering, consistent with a hyperfusion phenotype. Co-expression of Mfn1 or Mfn2 did not further enhance mitochondrial elongation in this setting. In contrast, in Mfn1/2 double knockout (DKO) MEF (**Fig. 5d**), FATE1 overexpression failed to induce mitochondrial elongation and instead cells presented with fragmented mitochondrial networks, similar to controls. While co-expression of Mfn1 and FATE1 in DKO cells led to perinuclear clustering of fragmented mitochondria, Mfn2 co-expression fully rescued the hyperfused phenotype. These results indicate that Mfn2, but not Mfn1, is both necessary and sufficient to mediate FATE1-driven mitochondrial elongation.

Together, these findings identify Mfn2 as a critical effector of FATE1-mediated mitochondrial hyperfusion and highlight a novel mechanism whereby FATE1 stabilizes mitochondrial networks by engaging Mfn2.

### FATE1 counteracts Drp1 recruitment and preserves mitochondrial network integrity under stress

Given FATE1’s lack of Drp1 recruitment capacity and its ability to promote mitochondrial hyperfusion under basal conditions, we next asked whether FATE1 expression could antagonize Drp1-mediated mitochondrial remodeling during stress-induced apoptosis. We tested three distinct stressors, namely TNF-α, valinomycin, and interferon-γ (IFN-γ), all known to trigger mitochondrial fragmentation through Drp1 activation (37, 38).

To assess Drp1 localization dynamics under these conditions, we co-expressed RFP-Drp1 with either OMM-GFP or GFP-FATE1 in HeLa cells and monitored Drp1 recruitment following acute stress exposure. In control cells, each treatment induced robust mitochondrial relocalization of RFP-Drp1 at 30 min, as visualized by the appearance of punctate mitochondrial Drp1 foci (**Fig. 6a**). In contrast, cells expressing GFP-FATE1 displayed significantly reduced Drp1 recruitment across all treatments, indicating that FATE1 can antagonize stress-induced Drp1 assembly at the mitochondrial surface. Quantification confirmed a consistent decrease in the percentage of cells positive for mitochondrial Drp1 in the presence of GFP-FATE1 (**Fig. 6c**).

**Figure 6:**
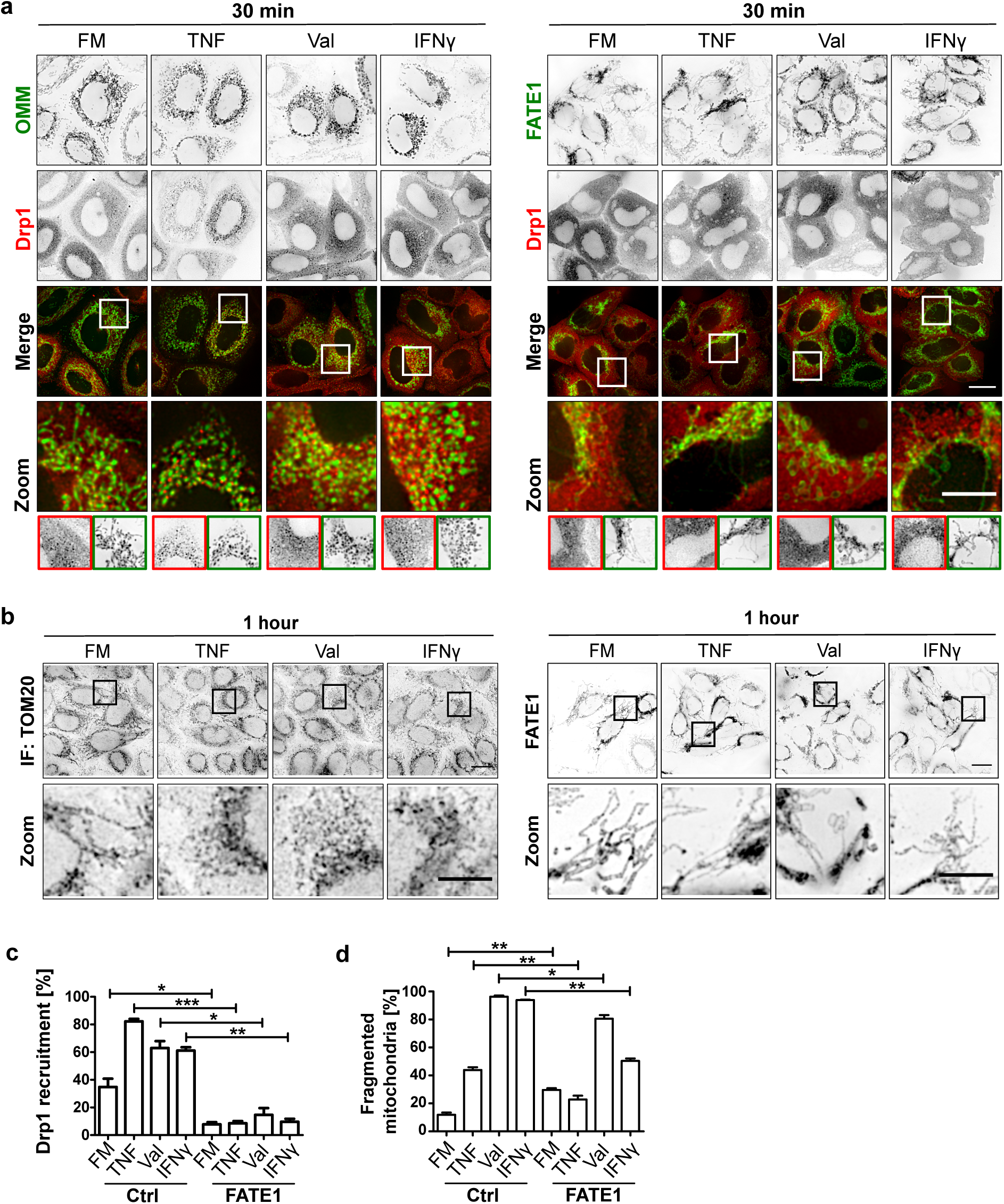
FATE1 counteracts Drp1 relocation and stabilizes mitochondrial networks upon cytokine- and mitochondrial uncoupling-induced stress. **a** HeLa cells were co-transfected with RFP-Drp1 and either OMM-GFP or GFP-FATE1. At 24 h post-transfection, cells were treated with TNF-α, valinomycin (Val) or IFN-γ for 30 min in full medium (FM), then fixed and imaged by widefield fluorescence microscopy. **b** HeLa cells transfected with GFP-Ctrl (top panels) or GFP-FATE1 (bottom panel) were treated as in (**a**), but for 1 h. Cells were fixed and IF-stained for Tom20 to visualize mitochondria in GFP-Ctrl expressing cells by widefield fluorescence microscopy. **c** Quantification of Drp1 mitochondrial recruitment from (**a**), scored as the percentage of cells showing Drp1 puncta on mitochondria. For each condition, 180 cells were analyzed in each of three independent experiments. Statistical comparisons to GFP-Ctrl were made per treatment using paired Student’s t-test; \**p*<0.05, \*\**p*<0.01, \*\*\**p*<0.001. **d** Quantification of mitochondrial fragmentation from (**b**), expressed as the percentage of cells with fragmented mitochondrial morphology. For each condition, 180 cells were analyzed in each of three independent experiments. Statistical comparisons to GFP-Ctrl were made per treatment using paired Student’s t-test; \*\**p*<0.01. Scale bars: overview, 10 µm; zoom, 5 µm.

We next examined whether this suppression of Drp1 recruitment correlated with protection against mitochondrial fragmentation. We expressed GFP-Ctrl or GFP-FATE1 in HeLa cells, followed by treatment with TNF-α, valinomycin, and IFN-γ for 1 h. While control cells exhibited pronounced fragmentation in response to all stressors, FATE1-expressing cells retained an elongated mitochondrial network (**Fig. 6b**). Morphometric quantification confirmed that FATE1 significantly reduced the proportion of cells with fragmented mitochondria across all treatment conditions (**Fig. 6d**).

Together, these findings demonstrate that FATE1 confers protection against stress-induced mitochondrial fragmentation by limiting Drp1 recruitment and preserving mitochondrial network integrity. These effects are consistent with FATE1’s role as a novel modulator of mitochondrial dynamics that opposes fragmentation and promotes hyperfusion under both basal and stress conditions.

## DISCUSSION

Mitochondrial dynamics, governed by a balance of fusion and fission events, are fundamental for maintaining organelle function, stress adaptation and cell fate decisions. Dysregulation of these processes is implicated in numerous pathologies, including cancer. In this study, we identify FATE1 as a novel modulator of mitochondrial morphology that antagonizes mitochondrial fission and promotes a hyperfused mitochondrial state, thereby enhancing resistance to stress-induced fragmentation of mitochondrial networks.

Although FATE1 shares structural similarity with the canonical fission adaptor Mff, including conserved C-terminal coiled-coil and transmembrane domains, it exhibits distinct subcellular localization and function. Our study identifies FATE1 as a unique modulator of mitochondrial morphology, acting independently of the fission factor Drp1 and instead promoting mitochondrial elongation through a mechanism dependent on the fusion protein Mfn2. Mff induces mitochondrial fission through recruitment of Drp1 to the outer mitochondrial membrane, a process mediated by two conserved N-terminal repeat motifs (R1 and R2) that are essential for Drp1 binding and oligomerization (7, 8, 31). Despite harboring a motif homologous to the R2 region of Mff, FATE1 lacks the R1 motif and does not promote Drp1 accumulation at mitochondrial membranes. This functional divergence is consistent with its dual localization to both mitochondria and the ER, in contrast to the strictly mitochondrial Mff.

Using domain-truncation mutants, and consistent with previous reports (20, 22), we show that the transmembrane domain of FATE1 is required for membrane integration, while the coiled-coil domain is essential for mitochondrial targeting. Interestingly, deletion of N-terminal segments enhanced mitochondrial localization, suggesting that this region may act as a negative regulator of mitochondrial insertion or promote ER retention. This points to a potential layer of regulatory control, whereby post-translational modifications or context-specific binding partners modulate FATE1 localization in response to cellular state. Future studies exploring such regulatory inputs may clarify how FATE1 activity is tuned under basal versus stress conditions, or across different tumor contexts.

Functionally, FATE1 overexpression induces a spectrum of mitochondrial phenotypes with a predominant shift toward hyperfusion, including a distinctive perinuclear clustering pattern. The hyperfused mitochondrial network observed in FATE1-expressing cells is reminiscent of stress-induced mitochondrial elongation, a protective adaptation against apoptotic and metabolic insults (39, 40). Mechanistically, we find that FATE1 selectively interacts with Mfn2, and that Mfn2 is both necessary and sufficient to mediate FATE1-driven mitochondrial elongation. In Mfn1/2 double-knockout MEFs, FATE1 fails to restore elongated mitochondria unless Mfn2 is co-expressed, pinpointing Mfn2 as a critical effector.

These mechanistic insights distinguish FATE1 from Mff, and highlight FATE1 as a functionally noncanonical modulator of mitochondrial dynamics. Its dual localization to the OMM and ER, along with its interaction with Mfn2, aligns with prior reports of FATE1 enrichment at mitochondria-associated membranes (MAMs), a key subdomain implicated in ER-mitochondria tethering and calcium signaling (22, 41). Through this positioning, FATE1 may simultaneously influence mitochondrial morphology and inter-organelle communication, with potential implications for bioenergetic adaptation and stress signaling in cancer cells.

Importantly, we found that the pro-fusion activity of FATE1 protects mitochondria from stress-induced mitochondrial fragmentation. FATE1 overexpression preserves elongated networks under cytotoxic stress, whereas its knockdown promotes mitochondrial fission, consistent with an earlier observation in adrenocortical carcinoma cells (22). We propose that FATE1 enforces a hyperfused mitochondrial state to maintain organelle integrity and suppress apoptosis initiation under cytotoxic stress.

Given its restricted expression profile and its role in suppressing stress-induced mitochondrial fragmentation and apoptosis, FATE1 represents a potential therapeutic target in tumors where mitochondrial hyperfusion supports survival. Recent analyses correlating elevated FATE1 expression with poor prognosis in cancers such as adrenocortical carcinoma (22, 42), colorectal cancer (20) and breast cancer (43) further underscore its clinical relevance. Future studies exploring pharmacologic modulation of FATE1 or its interaction partners may yield novel approaches for sensitizing tumor cells to apoptosis-inducing therapies.

Collectively, our work delineates a previously uncharacterized pathway through which FATE1 controls mitochondrial morphology and cell fate decisions, positioning it as a key modulator of mitochondrial stress responses with implications for cancer cell survival and response to therapeutic interventions.

## MATERIALS AND METHODS

### Cell culture and transient transfections

Human MCF-7 (CLS Cell Lines Service, Cytion, DE; no. 300273) breast cancer, and HeLa (CLS Cell Lines Service, Cytion: no. 300194) cervical cancer cell lines were maintained in DMEM (1g/L D-glucose, 0.11 g/L sodium pyruvate, Invitrogen). Human embryonic kidney 293T (CLS Cell Lines Service, Cytion; no. 300189), and wild type and Mfn1/Mfn2 double knockout (DKO) mouse embryonic fibroblasts (MEFs; ATCC; cat.no. CRL-2994) were maintained in DMEM (4.5 g/L D-glucose, Invitrogen). Media were supplemented with 10% FBS (Sigma Aldrich), L-glutamine (Gibco), non-essential amino acids (Gibco), and penicillin, streptomycin, amphotericin B (Gibco).

### Lentivirus-mediated gene knockdown

For lentiviral gene knockdown, human FATE1 targeting sequences (1_3: 5’-CAGCCAAACGAGTTTGGAATA-3’; 2: 5’-GAAATGATGGAGCTTGGATCT-3’; 3_2: 5’-TTGTGGAAAGGACGAAACA-3’) and a non-targeting control (5’-AATTGCCAGCTGGTT CCATCA-3’) were cloned into the 3rd generation lentiviral shRNA backbone pLKO.1 vector (Addgene #8453, RRID:Addgene_8453) (44)). HEK293T cells were transfected with pLKO.1 shRNA vectors together with the packaging vector psPAX2 (Addgene #12260, RRID:Addgene_12260) and the VSV-G envelope protein expression vector pMD2.G (Addgene #12259, RRID:Addgene_12259) using calcium-phosphate (Sigma Aldrich) transfection kits. Virus particle-containing supernatants were harvested and used to infect HeLa cells. Stable cell lines were selected and maintained with puromycin (1 µg/mL; Sigma Aldrich). Cells were plated and kept in puromycin-free medium at least 24 h prior experiments.

### Real-time PCR

Knockdown efficiency of FATE1-specific shRNAs was quantified by real-time quantitative PCR (qPCR). Total RNA was extracted using the RNeasy kit (Qiagen), and 1µg of RNA was reverse transcribed into cDNA using GoScript Reverse Transcription Mix, Oligo(dT) (Promega). HeLa cDNA samples were subjected to qPCR in triplicates using Platinum SYBR Green qPCR SuperMix (Invitrogen) and the StepOnePlus Real-Time PCR System (Applied Biosystems). For FATE1 forward 5’-CAAGAGCACCTGGTGATAGCAG-3’ and reverse 5’-ACTCGTTTGGCTGAAGCAGAGC-3’ primers (OriGene; cat.no. HP216304) were used, and for the reference housekeeping gene β-actin forward 5’-CACCATTGGCAATGAGCGGTTC-3’ and reverse 5’-AGGTCTTTGCGGATGTCCACGT-3’ (OriGene; cat.no. HP204660). Relative transcription levels were defined based on the expression levels of the target genes compared to the housekeeping gene using the 2^-ΔΔCT method (45).

### Gene expression and drug treatments

Transient transfections were performed using JetPRIME transfection reagent (Polyplus), according to the manufacturer’s instructions. Treatments with interferon-γ (IFN-γ; 1.1 µg/mL; German Cancer Research Center, Heidelberg, DE), tumor necrosis factor-α (TNF-α; 43 ng/mL; BASF), and valinomycin (VAL; 0.9 µM; Sigma) were performed in fully supplemented media (full media, FM).

### Plasmids

Fluorescent protein constructs generated for this study: Mfn1 (NM_033540.2) and SF-1 (NM_004959.4) cDNAs were obtained from DKFZ Core Facilities (German Cancer Research Center, Heidelberg, DE), PCR-amplified, and sub-cloned into either pmCherry-C1 or pEGFP-C1 or pmTagRFP-T-C1. Mfn2 was sub-cloned into pEGFP-C1 from Mfn2-YFP (Addgene, #28010; RRID:Addgene_28010 (9)). The ER retention sequence KDEL was cloned into ptagRFP-N1 to generate the ER marker RFP-KDEL.

FATE1 and Mff cDNAs were generated by reverse transcription PCR (RT-PCR) from total RNA isolated from HeLa and MCF-7 cells, respectively. PCR products were sub-cloned into pEGFP-C1, pmCherry-C1, or pcDNA3.1(−) to generate fluorescent protein-tagged and untagged (pcDNA3.1(−)), full-length (wild type, wt) FATE1 and Mff expression vectors. FATE1 truncation mutants were generated by PCR amplification from the full-length FATE1 cDNA using region-specific primer pairs incorporating restriction sites compatible with the expression vector (Sigma Aldrich). The Δ1-52, Δ1-130, Δ1-163, Δ89-184 and Δ163-184 (ΔTM) truncation mutants were generated by standard one-step PCR using forward and/or reverse primers specific to the desired truncation boundaries in combination with full-length reverse and/or forward primers, respectively. The resulting PCR products were gel-purified and subcloned into pEGFP-C1 to generate GFP-tagged FATE1 mutants. The ΔCC mutant was generated using a two-step overlap extension PCR strategy. In the first round, two separate PCRs were performed using the full-length forward primer and an internal reverse primer and the full-length reverse primer together with an internal forward primer. The resulting fragments were gel-purified, mixed in equimolar amounts, and used as template for a second PCR using the full-length forward and reverse primers to amplify the complete ΔCC delection mutant. All constructs were verified by Sanger sequencing (GATC Biotech).

Fluorescent protein constructs encoding fusion proteins previously described: mCherry-Drp1 wild type (34), dominant negative (DN) mCherry-Drp1(K38A) mutant (34), mito-mCherry (46) and GFP-OMM (47). Throughout this study, mCherry and tagRFP are referred to as RFP, and pEGFP as GFP.

### High-resolution fluorescence microscopy

Cells were plated in 8-well microscopy µ-slides (Ibidi; no. 80826). Transfections were performed as indicated, followed by fixation with 4% electron microscopy grade, methanol-fee paraformaldehyde (PFA; Alfa Aesar) in PBS (pH 7.4) for 15 min at room temperature. Cells were permeabilized using 0.3% Triton X-100 (Sigma Aldrich) in PBS for 10 min and blocked with 3% BSA (Sigma Aldrich) in Triton X-100/PBS for 45 min at room temperature. For immunofluorescence, cells were then incubated with primary antibodies against Tom20 (Santa Cruz Biotechnology, no. sc-11415 or sc-17764; 1:200) for 1 h at room temperature, or against COX IV (Cell Signaling, no. 4850S; 1:250), or TRAP1 (Novus Biologicals, no. NB300-555; 1:100) antibodies at 4°C overnight. Highly cross-absorbed Alexa Fluor 488, Alexa Fluor 546 and Alexa Fluor 647 secondary antibodies (Invitrogen) were used for immunofluorescence staining for 30 min at room temperature. Following primary and secondary antibody incubations, cells were washed three times in PBS. For visualization of mitochondria in live cells, cells were stained with 100 nM MitoTracker (MitoTracker Red FM, Invitrogen) for 15 min at 37 °C.

Fluorescence microscopy was performed using DeltaVision RT (Applied Precision) and DeltaVision Elite (GE Healthcare) widefield deconvolution systems equipped with 60x oil immersion objectives. Z-stacks were acquired with a step size of 0.3 µm and, for widefield microscopy, subsequently deconvolved using SoftWoRx software (Applied Precision). Image processing and analysis were carried out in Fiji/ImageJ (48). Representative images shown are maximum intensity projections of deconvolved Z-stacks. Single-channel images are displayed in inverted greyscale for optimal visibility of fine details (49). Font colors used in figure panels match the corresponding pseudocolors of fluorescent proteins or IF signals in merged images. Detection of endogenous proteins via IF is denoted by “IF” in single-channel labels to distinguish them from fluorescent protein fusions.

Cells were scored manually for mitochondrial morphology according to the following criteria: (i) Fragmented: punctate or short rod-like mitochondria scattered throughout the cytoplasm, (ii) Elongated: fused mitochondria with a filamentous appearance forming a partially connected network, and (iii) Hyperfused: highly interconnected and continuous mitochondrial network, often extending across the cytoplasm and spanning over the nucleus.

### Super-resolution fluorescence microscopy

For stimulated emission depletion (STED) microscopy, cells were plated on glass slides, transfected with pcDNA-FATE1 and at 24 h of expression fixed with 4% PFA in PBS for 15 min at room temperature, followed by 5 min with ice cold methanol and 2 min with ice cold acetone. Cells were then incubated with primary antibodies against Tom20 (Santa Cruz, no. sc-11415; 1:200) or FATE1 (Santa Cruz Biotechnology, no. sc-101220; 1:100) for 1 h at room temperature, or against COX IV (Cell Signaling, no. 4850S; 1:50) or TRAP1 (Novus Biologicals, no. NB300-555; 1:100) antibodies at 4°C overnight. Secondary antibodies anti-rabbit Atto565 (MPI, Department of Nanoptics, Goettingen, DE) and anti-mouse DY 485 XL (Dyomics) were incubated at room temperature for 45 min.

The two-color STED microscopy setup used for imaging was custom-built. Pulsed excitation was achieved using a high-repetition rate laser source PicoTA (PicoQuant) at a wavelength of λ=532 nm and as a second laser source LDH Series (PicoQuant) at a wavelength of λ=470 nm, synchronized with the STED laser triggered output of the laser. Fluorescence inhibition at λ=647 nm was accomplished by a rainbow laser (Spectra Physics). Beams were combined via an acousto-optical tunable filter (AOTF, Crystal Technologies) and coupled into a microscopy stand (DMI4000B, Leica Microsystems) equipped with a YANUS II laser scanner (Till Photonics) and an ACS APO, 63x/1.3 NA oil immersion lens (Leica Microsystems). AOTFs enable blanking of the lasers and allow controlling the laser power independently. Further they provide means for selecting the fluorescence returning from the microscope. The collected fluorescence was passed through an additional band-pass filter (580/40 (AHF Analysentechnik) and detected TAU-SPAD-100 (PicoQuant). A vortex phase plate (RPC Photonics) was used in the STED beam path to introduce a donut shaped focal intensity distribution.

### Western blotting

Whole cell lysates were prepared using RIPA buffer (Millipore) supplemented with PhosSTOP phosphatase and EDTA-free protease inhibitors (Roche). Equal amounts of protein per condition were electrophoresed on BisTris NuPAGE gels (Invitrogen) and transferred to nitrocellulose using the iBlot2 dry blotting system (Invitrogen). Subsequently, membranes were blocked for 2 h at room temperature and incubated with the following primary antibodies for 2 h at room temperature or overnight at 4°C: cytochrome *c* (BD Biosciences, no. 556433; 1:500), Drp1 (Cell Signaling, no. 8570S; 1:1000), COX IV (Cell Signaling, no. 4850S; 1:1000), GAPDH (Santa Cruz, no. sc-25778; 1:2000), GFP (Roche Applied Science, no. 11814460001), Mfn1 (Santa Cruz, no. sc-166644; 1:200), Mfn2 (Cell Signaling, no. 11925S; 1:1000), RFP (Chromotek, no. 5F8; 1:1000), Tim23 (BD, no. 611223; 1:3000), and VDAC (Cell Signaling, no. 4866; 1:1000). Following three washes, membranes were incubated with horseradish peroxidase-conjugated secondary, anti-mouse (GeneTex, no. GTX213111-01; 1:5000), anti-rabbit (GeneTex, no. GTX213110-01; 1:5000), or anti-rat (Cell Signaling, no. 7077S; 1:1000) antibodies for 1 h at room temperature. Following three washes, membranes were developed using SuperSignal West Pico (Thermo Fisher Scientific) chemiluminescent substrate and an Intas imager (Intas). Blots shown are representative of at least three independent experiments. Densitometric band quantifications were performed with ImageJ (50).

### Co-Immunoprecipitation

6 × 10^5^ cells were plated in 60-mm dishes on the day prior to transfections. At 24 h of expression, cells were lysed directly on ice in 150 µL CHAPS lysis buffer (2% CHAPS, 1% Triton X-100, 137 mM NaCl, 2 mM EDTA, 10% glycerol, and 20 mM Tris-HCl, pH 8.0) supplemented with protease and phosphatase inhibitors. A 5 µL aliquot of lysate was reserved as input, and the remaining 145 µL was incubated with 12.5 µL anti-GFP (1:40, Roche)-coupled Dynabeads Protein G (Invitrogen) prepared as follows: beads were first incubated with anti-GFP antibody for 15 min at room temperature with rotation, washed in 0.2M triethanolamine (Sigma Aldrich), and crosslinked using 20 mM dimethyl pimelimidate (DMP) in triethanolamine buffer for 30 min at room temperature. Crosslinking was quenched in 50 mM Tris buffer for 15 min at room temperature, and beads were washed in 0.02% PBS-T prior to use. Lysate-bead mixtures were rotated for 2 h at 4°C. Beads were washed with 0.02% Tween-20/PBS. Immunoprecipitated proteins were eluted by adding 2× LDS sample buffer and 1× sample reducing agent (both Invitrogen), followed by heating at 70°C for 10 min. Eluted, denatured proteins were analyzed by Western blotting using HRP-conjugated anti-mouse TrueBlot Ultra (Rockland, no. 18-8817-33; 1:1000) and anti-rabbit TrueBlot (Rockland, no. 18-8816-33; 1:1000).

### Mitochondrial isolation

Cells were seeded at 0.6 × 10^6^ cells/well of a 6-well plate and transfected the following day with GFP-Control, GFP-FATE1 or GFP-MFF plasmids. At 24 h post-transfection, cells were collected by scraping on ice and centrifuged at 500 × g for 5 min at 4°C. Cell pellets were resuspended in ice-cold mitochondrial isolation buffer (140 mM KCl, 10 mM MgCl_2_, 10 mM MOPS, 5 mM KH_2_PO_4_, 1 mM EGTA, 0.2% fatty acid-free BSA, pH 7.2) supplemented with EDTA-free protease inhibitor cocktail (Roche). Cells were disrupted using a ball-bearing cell homogenizer (Isobiotec) with a clearance of 8 µm and 10 strokes per sample on ice. Subcellular fractions were separated via differential centrifugation. Crude lysates were first centrifuged at 600 × g for 10 min at 4°C to remove nuclei and unbroken cells. The resulting supernatant was centrifuged at 2,800 × g for 10 min at 4°C to obtain a heavy membrane pellet enriched in mitochondria (P2). The supernatant was then subjected to centrifucation at 12,000 × g for 15 min at 4°C separate the rest-mitochondrial/light membrane fraction (P1) and cytosolic supernatant (S1). Pellets (P2 and P1) were resuspended in 50 µl 2× LDS sample buffer containing 1× reducing agent immediately after isolation. An equal volume of the cytosolic supernatant (S1) was also mixed with 2× LDS buffer. All fractions (P2, P1 and S1) were then subjected to SDS-PAGE followed by immunoblotting analysis. Successful fractionation was verified by marker immunoblotting: VDAC and Tim23 confirmed mitochondrial enrichment in P2; cytochrome *c* was primarily detected in P2 with minor presence in P1, consistent with intact membranes; and GAPDH, while predominantly cytosolic, showed detectable levels in P2 and P1, consistent with partial cytosolic contamination or known loose mitiochondrial association of glycolytic enzymes.

### Sequence analysis

Sequence alignments were performed using Jalview’s T-Coffee algorithm under default settings (51). COILS-based prediction algorithms were used to analyze the presence of a potential coiled-coil domain in FATE1 (29, 30). Mff served as a positive control and Bnip3 as a negative control.

### Statistical analysis

*p*-values were calculated with Student’s *t* test using GraphPad Prism for Windows (GraphPad Software, Boston, MA, USA; www.graphpad.com). Figure legends describe specific test settings. Unless stated otherwise, values are represented as mean ± S.E.M. of at least three independent experiments.

## DATA AVAILABILITY

All relevant data are within the paper and its supporting information files.

## ACKNOWLEDGEMENTS

We thank Dr. Stefan Hell (German Cancer Research Center and Bioquant, University of Heidelberg, Heidelberg, Germany) for use of his custom-built STED microscope. Mfn2-YFP was a gift from Richard Youle (Addgene plasmid # 28010). psPAX2 (Addgene plasmid # 12260) and pMD2.G were gifts from Didier Trono (Addgene plasmid # 12259), and pLKO.1 puro was a gift from Bob Weinberg (Addgene plasmid # 8453). This work was supported by the e:Bio grant #0316191 (LysoSys) of the Federal Ministry of Education and Research (BMBF), Germany (A.H.-B.), and funds #160650086 of the Department of Molecular Microbiology and Immunology, Johns Hopkins Bloomberg School of Public Health (A.H.-B.). The funders had no role in study design, data collection and analysis, decision to publish, or preparation of the manuscript. V.L. contributed to this study as a PhD candidate with A.H.-B. at the German Cancer Research Center (Heidelberg, Germany), and is currently affiliated with Monte Rosa Therapeutics Inc. (Basel, Switzerland). J.T.Z. is currently affiliated with Human Diagnostics Inc. (Wiesbaden, Germany), and N.R.B. with Humanity Neurotech Inc. (San Francisco, California, USA). No funding or support was received from Monte Rosa Therapeutics, Human Diagnostics, or Humanity Neurotech.

## AUTHOR CONTRIBUTIONS

Conceptualization, V.L., N.R.B., A.H.-B.; Methodology, V.L., J.T.Z., N.R.B., A.H-B.; Formal Analysis, V.L., J.T.Z., N.R.B., A.H-B.; Investigation, V.L., J.T.Z., D.P.T., N.R.B., A.H-B.; Data Curation, V.L., J.T.Z., N.R.B., A.H-B.; Writing – Original Draft, V.L., A.H.-B.; Writing – Review & Editing, J.T.Z., D.P.T., N.R.B.; Visualization, V.L., J.T.Z., D.P.T., A.H-B.; Supervision, N.R.B. and A.H.-B.; Funding Acquisition, A.H.-B.

## SUPPLEMENTARY MATERIAL

This article contains supplementary material.

