## Supplementary Figure 1 for "The Cancer/Testis Antigen FATE1 Antagonizes Fission and Preserves Mitochondrial Network Integrity under Cytotoxic Stress"

This PDF file includes:

Supplementary figure1 1

Legend for supplementary figure

### SUPPLEMENTARY FIGURES

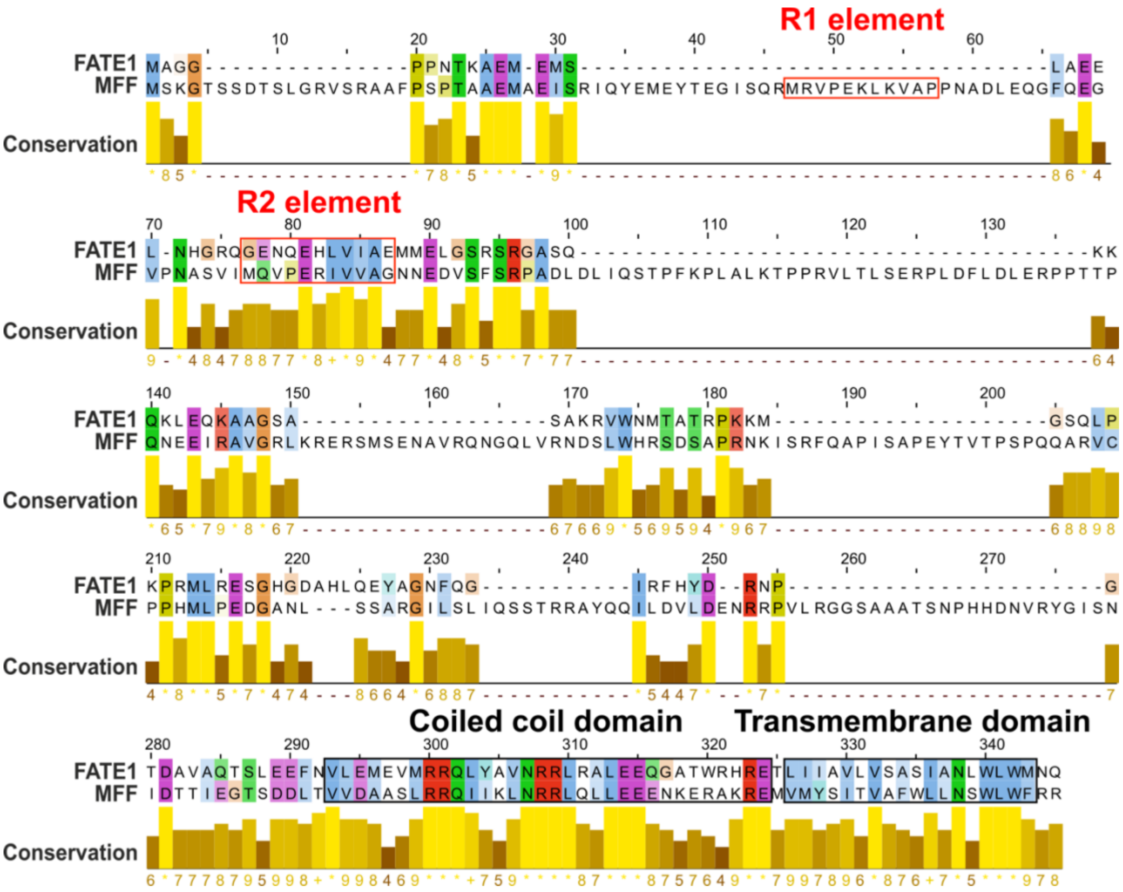


**Supplementary Fig. S1: Sequence alignment of human FATE1 and Mff.**

Sequence alignment of the C-terminal regions of human FATE1 and Mff was performed using Jalview with the T-Coffee algorithm under default parameters. Amino acid numbering corresponds to Mff. Red rectangles indicate the conserved R1 and R2 repeat motifs in Mff, which are essential for Drp1 recruitment and mitochondrial fission. FATE1 lacks a region homologous to the R1 motif and contains only a putative sequence resembling R2. Black rectangles denote the predicted coiled-coil (CC) and transmembrane (TM) domains in both proteins, with the CC and TM domains experimentally validated in Mff and putative homologous regions inferred in FATE1 based on sequence similarity.
